# Mpl expression on megakaryocytes and platelets is dispensable for thrombopoiesis but essential to prevent myeloproliferation

**DOI:** 10.1101/003459

**Authors:** Ashley P. Ng, Maria Kauppi, Donald Metcalf, Craig D. Hyland, Emma C. Josefsson, Marion Lebois, Jian-Guo Zhang, Tracey Baldwin, Ladina Di Rago, Douglas J. Hilton, Warren S. Alexander

## Abstract

Thrombopoietin (TPO) acting via its receptor Mpl is the major cytokine regulator of platelet number. To precisely define the role of specific hematopoietic cells in TPO dependent hematopoiesis, we generated mice that express the Mpl receptor normally on stem/progenitor cells but lack expression on megakaryocytes and platelets (*Mpl*^*PF4cre/PF4cre*^). *Mpl*^*PF4cre/PF4cre*^ mice displayed profound megakaryocytosis and thrombocytosis with a remarkable expansion of megakaryocyte-committed and multipotential progenitor cells, the latter displaying biological responses and a gene expression signature indicative of chronic TPO over-stimulation as the underlying causative mechanism, despite a normal circulating TPO level. Thus, TPO signaling in megakaryocytes is dispensable for platelet production; its key role in control of platelet number is via generation and stimulation of the bipotential megakaryocyte precursors. Nevertheless, Mpl expression on megakaryocytes and platelets is essential to prevent megakaryocytosis and myeloproliferation by restricting the amount of TPO available to stimulate the production of megakaryocytes from the progenitor cell pool.

**Significance statement:** Blood platelets, the small circulating cells that co-ordinate hemostasis, are produced by specialized bone marrow cells called megakaryocytes. The cytokine thrombopoietin (TPO) is a key regulator of platelet production acting via its specific cell receptor, Mpl. Via genetic modification of the *Mpl* allele in mice, we precisely define the bone marrow cells that express Mpl and, by genetically removing Mpl from megakaryocytes and platelets, we show TPO signaling via Mpl is not required in megakaryocytes for their expansion, maturation or platelet production. Rather, Mpl expression on megakaryocytes is essential for regulating TPO availability in the bone marrow microenvironment to prevent myeloproliferation, a model we suggest is important for human disease.

## Introduction

Thrombopoietin (TPO) is the principal hematopoietic cytokine that regulates platelet production at steady state and is required for rapid responses to platelet loss. TPO acts by binding to a specific cell surface receptor, Mpl, leading to receptor dimerization, activation of intracellular signal transduction pathways and responses of target cells. Mice lacking TPO or Mpl are severely thrombocytopenic and deficient in megakaryocytes and their progenitor cells, a phenotype consistent with a role for TPO in maintaining appropriate megakaryocyte numbers in vivo. In addition to its role in megakaryopoiesis, TPO is also an indispensible regulator of hematopoietic stem cells (HSC), essential for maintenance of quiescence and self-renewal (1).

TPO is produced primarily in the liver (2) and upon binding to the Mpl receptor on target cells is internalised and degraded. The prevailing model posits that circulating TPO concentration is inversely proportional to the “Mpl-mass” contributed by the total number of megakaryocytes and platelets. In normal individuals this model describes an effective feedback system to regulate TPO-driven megakaryocyte and platelet production according to need. The reciprocal relationship between platelet number and circulating TPO level is clearly evident in bone marrow transplant patients (1), and the key role of the TPO receptor is illustrated by the elevated circulating TPO in *Mpl*^−/−^ mice (3) and the modest elevation of platelet counts in transgenic mice expressing low levels of Mpl (4, 5). However, the relationship between circulating TPO concentration and peripheral platelet counts is not always conserved in pathological states of thrombocytosis and thrombocytopenia (6–9) suggesting that a simple relationship between megakaryocyte and platelet “Mpl-mass”, circulating TPO concentration and the degree of stimulation of megakaryopoiesis may not always hold.

While expression of Mpl on megakaryocytes and platelets contributes to regulation of available TPO, the role of direct TPO stimulation of megakaryocytes for effective platelet production is unclear. Administration of TPO in vivo or stimulation of bone marrow in vitro elevates megakaryocyte numbers and increases mean DNA ploidy (10, 11) and the thrombocytopenia in *TPO*^−/−^ mice is accompanied by reduced megakaryocyte ploidy (12). However, while exposure of megakaryocytes to TPO stimulates intracellular signaling (13), in vitro studies suggesting direct action of TPO on megakaryocytes to increase DNA ploidy, promote cytoplasmic maturation or to stimulate proplatelet production (14–16) are balanced by reports that TPO is dispensable for these megakaryocyte functions (15, 17–19).

To comprehensively define Mpl-expressing stem and progenitor cell populations in vivo and resolve the specific requirements for Mpl expression in megakaryocytes and platelets for platelet production and feedback control of TPO levels, we generated a novel mouse strain in which green fluorescent protein (GFP) is expressed from the *Mpl* locus and, when crossed to *Platelet Factor 4(PF4)cre* transgenic mice (20), specifically lacks Mpl expression in megakaryocytes and platelets.

## Results

A targeting vector was constructed for generation, via homologous recombination in embryonic stem cells, of a modified *Mpl* allele (*Mpl*^*fl/fl*^) designed to retain *Mpl* expression and provide a green fluorescent protein (GFP) reporter for transcriptional activity of the *Mpl* locus. The inclusion of *loxP* sites allowed for cre recombinase-mediated deletion of the GFP reporter as well as exons 11 and 12, resulting in a null allele (Fig. 1A and *SI Materials and Methods*). *PF4-cre* transgenic mice (20) have been reported to allow specific and efficient deletion of conditional alleles in megakaryocytes and platelets. To verify the activity and cell-type specificity of *PF4-cre* mice, particularly in view of a recent report of activity in HSCs (21), we crossed *PF4-cre* mice to mice in which enhanced yellow fluorescent protein (EYFP) is expressed in a cre-dependent manner (*Rosa26EYFP* (22)). While, as expected, EYFP was not expressed in the absence of cre recombinase, in *Rosa26EYFP*^*PF4Cre*^ mice EYFP was expressed in megakaryocytes and platelets, consistent with previous reports (20) and not in erythroid, lymphoid or myeloid cells (“SI Appendix, Fig. S1A”). Megakaryocyte formation progresses from a series of TPO-responsive bipotential erythroid-megakaryocyte progenitors (23). EYFP was not expressed in Lineage-negative Sca-1^+^ Kit^+^ (LSK) bone marrow cells, the population containing stem and primitive multipotential progenitor cells, nor in any progenitor cell fraction tested, including the BEMP and CD150^+^CD9^hi^ megakaryocyte/erythroid-restricted bipotential megakaryocyte-erythroid progenitor population and CFU-E erythroid progenitor population (“SI Appendix, Table S1, Fig. S1B,C”). EYFP expression was observed in a proportion of cells in a previously uncharacterised CD150^+^FcγR^+^ population (Lin^−^Sca-1^−^Kit^+^CD150^+^IL7Rα^−^FcγRII/III^+^Endoglin^lo^CD9^hi^, “SI Appendix, Fig. S1B,C”) that comprised ∽ 0.01% of the bone marrow (“SI Appendix, Fig. S2A”) and which demonstrated potent erythro-megakaryocytic potential in vitro and in vivo (“SI Appendix, Fig. S2B,C”). We conclude that the CD150^+^FcγR^+^ fraction includes megakaryocyte precursors downstream from the previously characterised CD150^+^CD9^+^ fraction, revealed by the onset of *PF4* promoter activity that, given the lack of YFP expression in CFU-E in *Rosa26EYFP*^*PF4Cre*^ mice, may define a megakaryocyte-restricted precursor within the bipotential CD150^+^FcγR^+^ fraction. Thus cre-dependent recombination using the *PF4-cre* transgenic mouse was restricted to late megakaryocyte progenitors, megakaryocytes and platelets and absent in other progenitor cells and hematopoietic lineages.

**Fig. 1.**
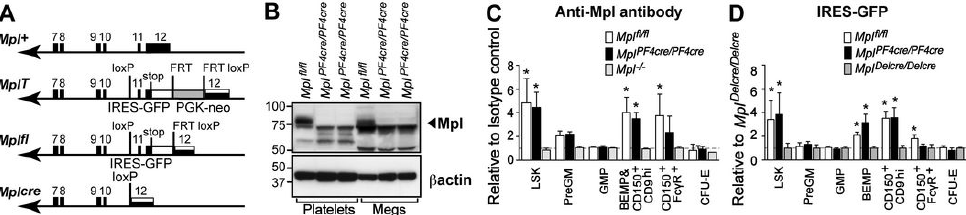
Targeted modification of the Mpl locus in mice and Mpl expression in *Mpl*^*PF4cre/PF4cre*^ mice. (A) Generation of a GFP reporter allele of *Mpl* expression that also allows conditional, cre-mediated *Mpl* inactivation. ***Mpl***^+^: wild type *Mpl* allele. Exons 7 – 12 (filled boxes), 3’ untranslated region of exon 12 (striped). ***Mpl^T^***: targeting vector incorporated into the *Mpl* locus. IRES-GFP cassette (white), PGK-neo selection cassette (shaded). ***Mpl*^*fl*^**; GFP reporter for expression from the *Mpl* locus. PGK-neo excised by intercrossing with Flp recombinase transgenic mice. ***Mpl^cre^***; *Mpl* null allele generated via cre-mediated excision. (B) Western blot of protein extracted from platelets (100 μg) and megakaryocytes (Megs, 188 μg) pooled from two to three independent genotype-matched *Mpl*^*fl/fl*^ and *Mpl*^*PF4cre/PF4cre*^ mice per lane. (C) The mean fluorescence intensity of Mpl expression on stem and progenitor cell populations from *Mpl*^*fl*^, *Mpl*^*PF4cre/PF4cre*^ and *Mpl*^−/−^ mice is shown relative to the isotype control (see “SI Appendix, Fig. S3C”). Mean and standard deviation shown, n=4 mice per genotype. * = *P* < 0.005 by two-tailed Student’s t-test. (D) Transcriptional activity of the *Mpl* locus via expression of the IRES-GFP cassette in stem and progenitor populations. GFP mean fluorescence intensity from *Mpl*^*fl/fl*^ and *Mpl*^*PF4cre/PF4cre*^ cells is shown relative to *Mpl*^*Delcre/Delcre*^ mice (see “SI Appendix, Fig. S3D”). Mean and standard deviation shown, n = 3 − 8 mice per genotype. * = *P* < 0.002.

### *Mpl* expression is absent in megakaryocytes and platelets, but retained in megakaryocyte progenitor and stem cells in Mpl ^PF4cre/PF4cre^ mice

To investigate the physiological effects of *Mpl* deletion on megakaryocytes and platelets we crossed *Mpl*^*fl/fl*^ mice to *PF4cre* mice to produce progeny in which the intracellular domain of *Mpl* and the IRES-GFP cassette had been specifically deleted in megakaryocytes and platelets (*Mpl*^*PF4cre/PF4cre*^). Western blot analysis of extracts from megakaryocytes and platelets demonstrated efficient ablation of Mpl expression in *Mpl*^*PF4cre/PF4cre*^ mice (Fig. 1B). This was confirmed by analysis of *Mpl-GFP* reporter expression: while megakaryocytes generated in culture from *Mpl*^*fl/fl*^ mice expressed GFP, this was lost in *Mpl*^*PF4cre/PF4cre*^ mice, consistent with recombination and deletion of the targetted *Mpl*^*fl*^ intracellular domain and IRES-GFP cassette in these cells (“SI Appendix, Fig. S3A”, Fig. 1A), and consequently, absence of GFP in platelets (“SI Appendix, Fig. S3B”). As anticipated, no activity of the *Mpl* locus was evident in lymphocytes, granulocytes or erythroid cells (“SI Appendix, Fig. S3B”). In LSK cells, *Mpl*^*PF4cre/PF4cre*^ mice expressed Mpl at equivalent levels to *Mpl*^*fl/fl*^ controls, both via flow cytometry with anti-Mpl antibody, and via the *Mpl-GFP* reporter (Fig. 1C,D and “SI Appendix, Fig. S3C,D”). As expected, *Mpl* expression was not detected in pre-granulocyte-macrophage (preGM) or granulocyte-macrophage (GMP) progenitor cells, nor in CFU-E. In *Mpl*^*fl/fl*^ control megakaryocyte progenitor cells, Mpl expression first became apparent in BEMP and was maintained in maturing CD150^+^CD9^hi^ and CD150^+^FcγR^+^ progenitor cells. As expected Mpl was not detected in any cell populations examined in previously generated *Mpl*^−/−^ mice (24), nor via the *Mpl-GFP* reporter in *Mpl*^*Delcre/Delcre*^ control mice, in which *Mpl*^*fl/fl*^ mice were crossed to *Deleter-cre* transgenic mice (25) to generate mice in which the *Mpl*^*fl*^ locus including the IRES-GFP reporter cassette were deleted throughout the animal. Importantly, Mpl expression was unperturbed in *Mpl*^*PF4cre/PF4cre*^ BEMP and CD150^+^CD9^hi^ megakaryocyte-biased bipotential progenitors, but reduction in *Mpl* IRES-GFP transcription and cell surface Mpl expression was apparent in *Mpl*^*PF4cre/PF4cre*^ CD150^+^FcγR^+^ megakaryocyte precursors.

Thus, in *Mpl*^*PF4cre/PF4cre*^ mice, Mpl expression was intact in LSK and early bipotential erythro-megakaryocytic progenitor cells, reduced in later progenitors with megakaryocyte potential and absent in megakaryocytes and platelets. The pattern of ablation of Mpl expression in *Mpl*^*PF4cre/PF4cre*^ mice was entirely consistent with the activity of *PF4-cre* defined in *Rosa26EYFP^PF4Cre^* mice.

### *Mpl*^*PF4cre/pF4cre*^ mice develop a marked thrombocytosis and megakaryocytosis

While *Mpl*^*PF4cre/PF4cre*^ mice were healthy and displayed no outward abnormalities, analysis of peripheral blood (Table 1) revealed a remarkable thrombocytosis with a 10-fold increase in the number of circulating platelets in comparison to *Mpl*^*fl/fl*^ or *Mpl*^+/+^ controls. In contrast, and as expected, *Mpl*^*Delcre/Delcre*^ exhibited a profound thrombocytopenia identical in phenotype to previously reported *Mpl*^−/−^ mice (24). In contrast, the numbers of other blood cells in *Mpl*^*PF4cre/PF4cre*^ mice were not significantly different from *Mpl*^*fl/fl*^ or *Mpl*^+/+^ controls. Significant splenomegaly was evident in *Mpl*^*PF4cre/PF4cre*^ mice (*Mpl*^*PF4cre/PF4cre*^ 210 ± 46 mg, *Mpl*^*fl/fl*^ 76±15 mg, n=4 per genotype, *P* = 0.0012) with histology revealing gross megakaryocytosis in the bone marrow and spleen, with the numbers of megakaryocytes in these organs even greater than observed in transgenic mice (*TPO*^*Tg*^, (3)) engineered to express high amounts of TPO (Fig. 2A, “SI Appendix, Fig. S4A”). Both *Mpl*^*PF4cre/PF4cre*^ and *TPO*^*Tg*^ mice had a significant increase in the proportion of megakaryocytes with ploidy of 16N or greater relative to *Mpl*^*fl/fl*^ controls (Fig. 2B). Clonogenic assays of bone marrow and spleen cells revealed a profound increase in the numbers of megakaryocyte colony-forming cells in *Mpl*^*PF4cre/PF4cre*^ mice (Table 2). The profound thrombocytosis in *Mpl*^*PF4cre/PF4cre*^ was not accompanied by significant extension in platelet lifespan (“SI Appendix, Fig. S4B”). Together, these data suggest that the thrombocytosis in *Mpl*^*PF4cre/PF4cre*^ mice is caused primarily by excess production of megakaryocytes and their progenitors.

**Table 1.**
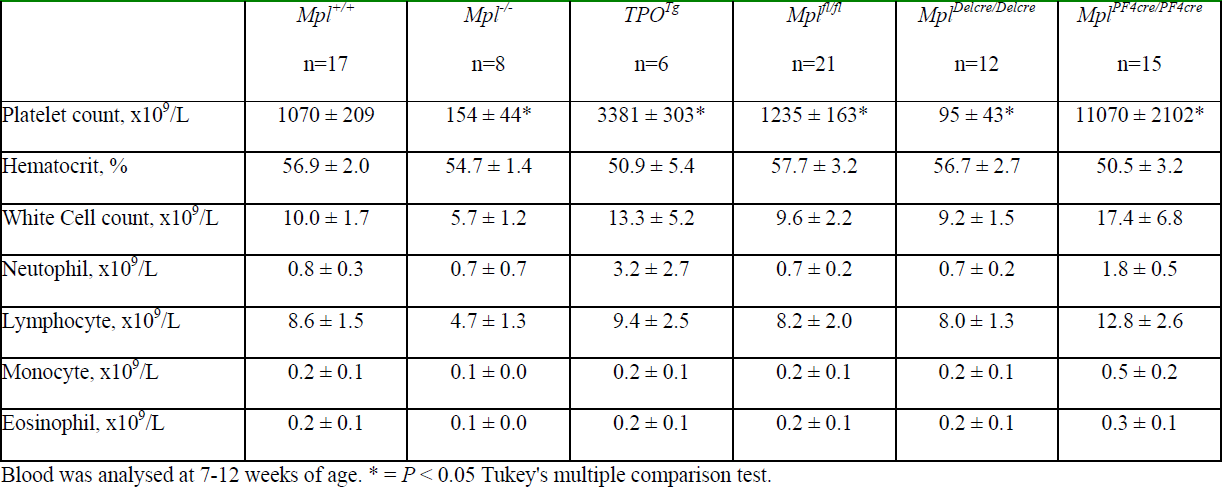
Peripheral blood cells in *Mpl*^*PF4cre/PF4cre*^ and control mice

**Fig. 2.**
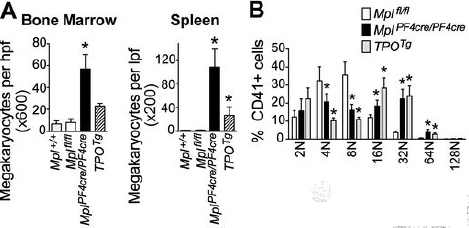
Expanded megakaryopoiesis in *Mpl*^*PF4cre/PF4cre*^ mice. (A) Number of megakaryocytes per high powered field from bone marrow (left) and spleen (right) of *Mpl*^+/+^, *Mpl*^*fl/fl*^, *Mpl*^*PF4cre/PF4cre*^ and *TPO*^*Tg*^ mice. * = *P* < 0.03 by Student’s unpaired two-tailed t-test, n = 3 − 9 mice per genotype. (B) Ploidy of bone marrow megakaryocytes from *Mpl*^*fl/fl*^, *Mpl*^*PF4cre/PF4cre*^ and *TPO*^*Tg*^ mice. Mean and standard deviation shown, n = 4-6 mice per genotype. * = *P*_adj_ < 0.03 by Student’s unpaired two-tailed t-test using Bonferroni testing for multiple comparisons.

### *Mpl*^*PF4cre/pF4cre*^ mice have perturbed stem and progenitor cell compartments and develop features of myeloproliferation

In addition to excess numbers of megakaryocyte progenitor cells, the total number of myeloid progenitor cells in the bone marrow and spleen of *Mpl*^*PF4cre/PF4cre*^ mice was also significantly elevated, with a particular increase in the number of blast colony-forming cells, a phenotype also observed in the bone marrow of *TPO*^*Tg*^ mice (Table 2). HSC and myeloid progenitor cell populations were analysed by flow cytometry, with particular emphasis on cells with megakaryocyte potential. *Mpl*^*PF4cre/PF4cre*^ mice displayed a two- to four-fold increase in bone marrow LSK cells, Lin^−^cKit^+^Scal^−^ myeloid progenitors, pre-granulocyte-macrophage (PreGM) progenitors, GM progenitors (GMP) and CD150^+^CD9^hi^ and CD150^+^FcγR^+^ bipotential erythroid-megakaryocyte progenitor populations, with a reduction in erythroid progenitors (preCFU-E and CFU-E) when compared to *Mpl*^+/+^ and/or *Mpl*^*fl/fl*^ controls (Fig. 3, “SI Appendix, Fig. S5”). This was similar to the effects caused by chronic excessive in vivo TPO stimulation (*TPO*^*Tg*^, Fig. 3, “SI Appendix, Fig. S5” and (23)), but was significantly more marked in *Mpl*^*PF4cre/PF4cre*^ mice.

**Table 2.**
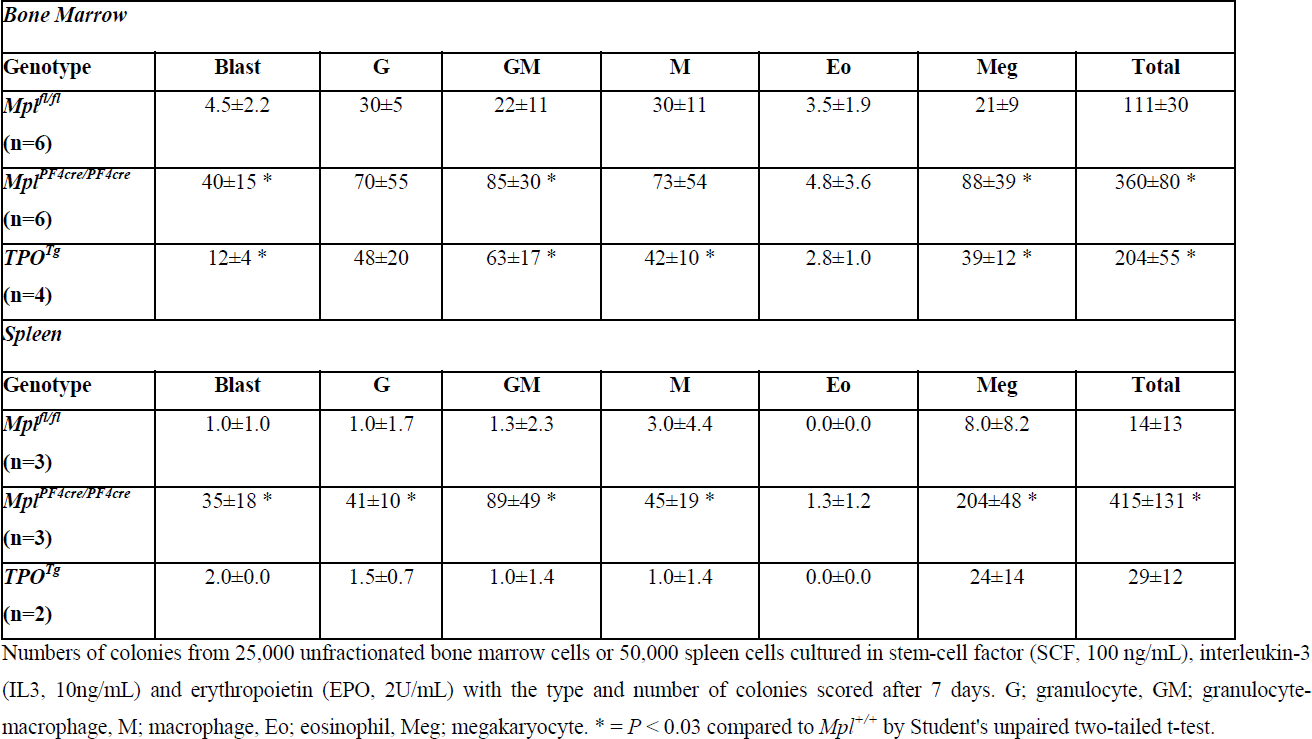
Numbers of clonogenic hemopoietic progenitor cells in *Mpl*^*PF4cre/PF4cre*^ and control mice

| **Bone Marrow** |  |  |  |  |  |  |  |
| --- | --- | --- | --- | --- | --- | --- | --- |
| **Genotype** | **Blast** | **G** | **GM** | **M** | **Eo** | **Meg** | **Total** |
| *Mpl*<sup>fl/fl</sup><br>(n=6) | 4.5±2.2 | 30±5 | 22±11 | 30±11 | 3.5±1.9 | 21±9 | 111±30 |
| *Mpl*<sup>PF4cre/PF4cre</sup><br>(n=6) | 40±15 * | 70±55 | 85±30 * | 73±54 | 4.8±3.6 | 88±39 * | 360±80 * |
| *TPO*<sup>Tg</sup><br>(n=4) | 12±4 * | 48±20 | 63±17 * | 42±10 * | 2.8±1.0 | 39±12 * | 204±55 * |
| **Spleen** |  |  |  |  |  |  |  |
| **Genotype** | **Blast** | **G** | **GM** | **M** | **Eo** | **Meg** | **Total** |
| *Mpl*<sup>fl/fl</sup><br>(n=3) | 1.0±1.0 | 1.0±1.7 | 1.3±2.3 | 3.0±4.4 | 0.0±0.0 | 8.0±8.2 | 14±13 |
| *Mpl*<sup>PF4cre/PF4cre</sup><br>(n=3) | 35±18 * | 41±10 * | 89±49 * | 45±19 * | 1.3±1.2 | 204±48 * | 415±131 * |
| *TPO*<sup>Tg</sup><br>(n=2) | 2.0±0.0 | 1.5±0.7 | 1.0±1.4 | 1.0±1.4 | 0.0±0.0 | 24±14 | 29±12 |

Numbers of colonies from 25,000 unfractionated bone marrow cells or 50,000 spleen cells cultured in stem-cell factor (SCF, 100 ng/mL), interleukin-3 (IL3, 10ng/mL) and erythropoietin (EPO, 2U/mL) with the type and number of colonies scored after 7 days. G; granulocyte, GM; granulocyte-macrophage, M; macrophage, Eo; eosinophil, Meg; megakaryocyte. \* = $P < 0.03$ compared to *Mpl*<sup>+/+</sup> by Student's unpaired two-tailed t-test.

**Fig. 3.**
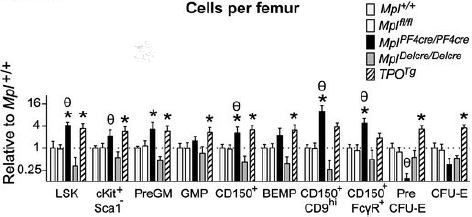
Expanded progenitors in *Mpl*^*PF4cre/PF4cre*^ and *TPO*^*Tg*^ mice. Numbers of cells in flow cytometrically-defined bone marrow stem and progenitor cell fractions from *Mpl*^*fl/fl*^ (n=8), *Mpl*^*PF4cre/PF4cre*^ (n=6), *Mpl*^*Delcre/Delcre*^ (n=3) and *TPO*^*Tg*^ (n=5) mice shown as cells per femur relative to *Mpl*^+/+^ (n=4) controls. For definitions of cell populations, see “SI Appendix, Fig. S5, Table S1”. Mean and standard deviation shown. * = *P*_adj_ < 0.05 by two-tailed Student’s unpaired t-test compared to *Mpl*^+/+^; θ = *P*_adj_ < 0.05 by two-tailed Student’s unpaired t-test specifically for *Mp*^*PF4cre/PF4cre*^ compared to *Mpl*^*fl/fl*^ with Bonferroni testing for multiple comparisons.

Blast cell colonies, which represent primitive multi-potential pre-progenitor cells (26) predominantly derived from the LSK population (27, 28) were picked from primary cultures and replated. Blast cell colonies from *Mpl*^*PF4cre/PF4cre*^ mice had a sigificantly increased propensity to formation of secondary colonies, an effect also evident in *TPO*^*Tg*^ mice, particularly secondary blast cell and megakaryocyte colonies (“SI Appendix, Fig. S6”), demonstrating a greater in vitro capacity for pre-progenitor cell self-renewal and secondary colony formation when compared to *Mpl*^*fl/fl*^ controls. This provides evidence at the single cell level that *Mpl*^*PF4cre/PF4cre*^ multipotential pre-progenitor cells, like those in *TPO*^*Tg*^ mice, have characteristics of chronic excessive TPO exposure.

### Stem and progenitor cells expressing Mpl are responsible for TPO clearance in *Mpl^PF4cre/PF4cre^* mice

Circulating TPO concentrations were assayed to determine how serum TPO levels were affected by loss of Mpl expression on megakarycytes and platelets in *Mpl*^*PF4cre/PF4cre*^ mice. As expected, *TPO*^*Tg*^ mice, engineered to express excess TPO, and *Mpl*^−/−^ mice, which have no capacity to clear serum TPO by Mpl receptor-mediated endocytosis, had elevated levels of serum TPO (Fig. 4A). Surprisingly, the circulating TPO level in *Mpl*^*PF4cre/PF4cre*^ mice, was not significantly different from *Mpl*^*fl/fl*^ controls. Transcription of TPO in the liver, the major site of TPO production, was unaltered in *Mpl*^*PF4cre/PF4cre*^ mice (Fig. 4B). This finding implies that Mpl-expressing LSK and megakaryocyte/erythroid progenitor cells in *Mpl*^*PF4cre/PF4cre*^ mice, which are expanded significantly in number, clear circulating TPO to levels observed in Mpl-replete mice, even in the absence of clearance by megakaryocytes and platelets.

**Fig. 4.**
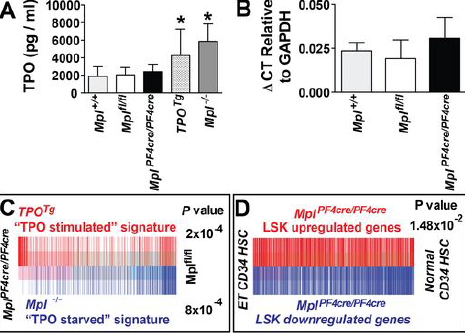
TPO transcription, circulating TPO concentration and gene expression changes in LSK cells of *Mpl*^*PF4cre/PF4cre*^ mice. (A) TPO concentration by immunoassay in serum from *Mpl*^+/+^, *Mpl*^*fl/fl*^, *Mpl*^*PF4cre/PF4cre*^, *Mpl*^−/−^ and *TPO*^*Tg*^ mice. * = *P* < 0.04, n = 10-17 mice per genotype. (B) TPO expression in livers determined by quantitative RT-PCR with ΔCT shown as mean and standard deviation relative to GAPDH expression. There were no significant differences between *Mpl*^+/+^, *Mpl*^*fl/fl*^, *Mpl*^*PF4cre/PF4cre*^ genotypes, n= 3-5 mice per group. (C) Barcode plot showing ability of *TPO*^*Tg*^ “TPO stimulated” LSK signature genes and *Mpl*^−/−^ “TPO starved” signature genes (see “SI Appendix, Table S3, S4”) to distinguish between *Mpl*^*PF4cre/PF4cre*^ LSKs and *Mpl*^*fl/fl*^ LSKs, with corresponding *P* values derived from rotational gene set testing using ROAST. Red bars designate upregulated genes in the *TPO*^*Tg*^ “TPO stimulated” LSK signature set (*P* = 2×10^−4^); blue bars designate *Mpl*^*−/−*^ “TPO starved” LSK signature set (*P* = 8×10^−4^); demonstrating gene expression in *Mpl*^*PF4cre/PF4cre*^ LSKs is strongly correlated with a “TPO stimulated” gene signature. (D) Barcode plot showing the TPO stimulated gene expression signature of *Mpl*^*PF4cre/PF4cre*^ LSKs (Red bars, upregulated genes; Blue bars, downregulated genes, see “SI Appendix, Table S2”) can distinguish between CD34^+^ bone marrow cells of patients with essential thromboocythemia from controls ((33) GEO database GSE9827, *P* = 1.48×10^−2^). *P* value derived from rotational gene set testing using ROAST with the *Mpl*^*PF4cre/PF4cre*^ LSK gene signature weighted by log fold-change (see “SI Appendix, Table S2, see also Fig. S7 and SI Materials and Methods”).

### Gene expression profiling of *Mpl*^*PF4cre/PF4cre*^ stem and primitive progenitors demonstrate a TPO stimulation signature

Gene expression profiling was undertaken on sorted LSK populations from *Mpl*^+/+^, *TPO*^*Tg*^, *Mpl*^−/−^, *Mpl*^*fl/fl*^ and *Mpl*^*PF4cre/PF4cre*^ mice using Illumina WG version 2 bead-chip microarrays. Pair-wise comparison between *Mpl*^*PF4cre/PF4cre*^ and *Mpl*^*fl/fl*^ LSKs was performed to obtain differentially expressed genes using a linear modeling and an empirical Bayes approach ((29), “SI Appendix, Table S2, and *SI Materials and Methods*”). Signature genes of *TPO*^*Tg*^ LSKs and *Mpl*^−/−^ LSKs were defined as those that were significantly differentially expressed in a positive direction by pair-wise comparison to wild-type LSKs (“SI Appendix, Table S3, S4”). Rotational gene set tests were then performed using ROAST (30) with the “TPO over-stimulation” gene signature from *TPO*^*Tg*^ LSKs and the “TPO starvation” gene signature from *Mpl*^−/−^ LSKs. This analysis revealed that genes associated with TPO overstimulation in *TPO*^*Tg*^ LSKs were strongly correlated with upregulated genes in *Mpl*^*PF4cre/PF4cre*^ LSKs whereas the gene signature associated with TPO starvation in *Mpl*^−/−^ LSKs was associated with downregulated genes in *Mpl*^*PF4cre/PF4cre*^ LSKs (Fig. 4C). This data confirms at a genetic level that Mpl expressing stem and progenitor cells in *Mpl*^*PF4cre/PF4cre*^ mice were exposed to excessive TPO stimulation, explaining the behaviour of their pre-progenitor blast colonies at the single cell level and the mechanistic basis for the myeloproliferation, megakaryocytosis and thrombocytosis obverved in *Mpl*^*PF4cre/PF4cre*^ mice.

### Human CD34^+^ stem and progenitor cells demonstrate a Mpl^PF4cre/PF4cre^ LSK gene expression signature

Megakaryocytes and platelets in human myeloproliferative neoplasms have been shown to express reduced levels of MPL (31, 32). To explore the possibility that reduced MPL expression on megakaryocytes and platelets in human disease might contribute to the myeloproliferative phenotype via a mechanism analagous to the excessive TPO stimulation of stem and progenitor cells evident in the *Mpl*^*PF4cre/PF4cre*^ model, we compared the gene expression signature of *Mpl*^*PF4cre/PF4cre*^ LSK cells (“SI Appendix, Table S2”) to gene expression in CD34^+^ bone marrow cells from patients with mutated *JAK2V617F* and *JAK2* wild-type essential thrombocythemia (GEO database GSE9827, (33)). Rotational gene set testing demonstrated significant enrichment of the *Mpl*^*PF4cre/PF4cre*^ LSK gene signature (*P* = 1.48×10^−2^, Fig. 4D) and *TPO*^*Tg*^ and Mpl^μ/μ^ LSK gene signatures (*P* = 8 × 10^−4^, “SI Appendix, Fig. S7”) in CD34^+^ cells from patients with disease. Thus, reduced expression of Mpl in megakaryocytes and platelets resulting in the excessive TPO stimulation of stem and progenitor cells that characterises *Mpl*^*PF4cre/PF4cre*^ mice may in part underpin the pathogenesis of megakaryocytosis and thrombocytosis in human myeloproliferative disease.

## Discussion

While it has been established TPO signaling is required for maintaining steady state platelet numbers and response to crises requiring rapid platelet production, the precise cellular mechanisms by which TPO actions are achieved remained largely undefined. Our results definitively establish that 1) Mpl expression on megakaryocytes is dispensable for high level platelet production and megakaryocyte maturation, 2) the primary mechanism by which TPO signaling stimulates thrombopoiesis is via Mpl-expressing hematopoietic progenitors, and 3) Mpl receptor on megakaryocytes and platelets acts predominantly, if not solely to regulate, via internalization and degradation, the amount of TPO available to hematopoietic stem and progenitor cell populations (Fig. 5).

**Fig. 5.**
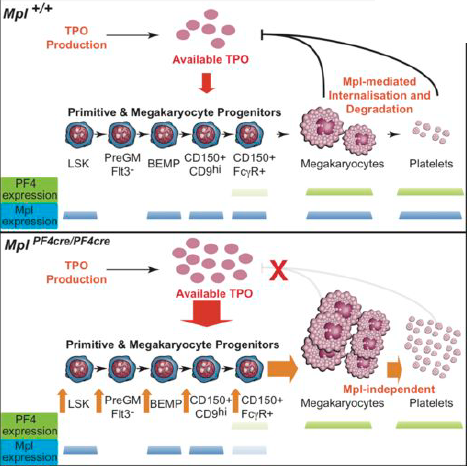
Model for regulation of TPO and control of megakaryopoiesis. (Top panel) Clearance of TPO by Mpl-expressing megakaryocytes in bone marrow and the peripheral blood platelet pool maintains TPO homeostasis at steady state and in situations of acute thrombocytopenia. For definitions of cell populations, see “SI Appendix, Table S1”. Mpl expression is shown as blue bars for each population in *Mpl*^*fl/fl*^ mice. (Bottom panel) Loss of TPO clearance by megakaryocytes and platelets leads to excessive TPO stimulation of Mpl-expressing HSCs and progenitor cells, multi-lineage progenitor expansion and differentiation towards the megakaryocyte lineage from bi-potential megakaryocyte-erythroid progenitors, with consequent myeloproliferation, megakaryocytosis and thrombocytosis. While the availability of TPO for stimulation of stem/progenitor cells is increased, consumption by the expanded numbers of these cells normalises circulating TPO concentration. PF4-cre expression is shown as green bars and Mpl expression is shown as blue bars for each population in *Mpl*^*PF4cre/PF4cre*^ mice.

Regarding the role of Mpl expression on megakaryocytes, while increased megakaryocyte ploidy has been associated with high TPO activity and the converse has been observed in TPO/Mpl-deficient mouse models (3, 10–12), a shift to higher ploidy megakaryocytes accompanied marked thrombocytosis in *Mpl*^*PF4cre/PF4cre*^ mice was observed despite the absence of Mpl expression in megakaryocytes. Thus, thrombopoiesis and the development of high DNA ploidy in megakaryocytes is independent of TPO signaling in these cells; the latter likely reflecting alternative mechanisms associated with states of increased megakaryocyte production. A role for TPO in sensitizing platelets to specific activators (34) may explain the retention of Mpl signaling pathways in megakaryocytes and platelets as our data do not exclude TPO, when functional Mpl is present, having a positive effect on megakaryocyte or platelet function.

Notably, *Mpl*^*PF4cre/PF4cre*^ mice exhibited all the phenotypic hallmarks and gene expression changes in stem/primitive multipotential progenitor cells, indicative of chronic excessive TPO stimulation. This was despite having normal concentrations of serum TPO. These data support a model in which absence of megakaryocyte and platelet Mpl mass allows greater available TPO for stimulation of Mpl-expressing stem/progenitor cells, the expansion of which normalises the circulating TPO concentration via Mpl-mediated internalisation. Thus, the circulating TPO concentration in this model is determined by the Mpl mass of both the megakaryocyte/platelet pool and Mpl-expressing stem and progenitor cells. Moreover, as *Mpl*^*PF4cre/PF4cre*^ mice were observed to have a more profound myeloproliferative phenotype with a greater degree of thrombocytosis relative to *TPO*^*Tg*^ counterparts, despite the latter having a higher circulating TPO level, suggests local TPO availability within the stem and progenitor cell microenvironment of the bone marrow may be regulated in significant part by megakaryocytes: *TPO*^*Tg*^ mice would retain megakaryocyte Mpl-mediated regulation of local TPO contentration, while absence of functional Mpl on megakaryocytes in *Mpl*^*PF4cre/PF4cre*^ mice, would significantly impair local TPO regulation. Increasingly, evidence has suggested a close physical relationship between megakaryocytes and bone marrow stem/progenitor cell niches. Imaging studies have identified megakaryocytes colocalising with CD150^+^CD48^−^Lin^−^ HSCs adjacent to sinusoidal endothelium (35) as well as in close proximity to Nestin^+^ mesenchymal stem cells, which form a significant functional component of the HSC niche (36).

The importance of TPO as a multi-functional regulator of hematopoiesis was highlighted. Chronic TPO overstimulation in *Mpl*^*PF4cre/PF4cre*^ and *TPO*^*Tg*^ mice resulted in excess numbers of primitive multipotential pre-progenitor blast colonies that are principally derived from LSK cells, a population that expresses Mpl. The increased propensity of pre-progenitor cells to self-renew and to produce increased numbers progenitors of multiple myeloid lineages with chronic TPO overstimulation provide a mechanistic rationale for the effectiveness of the TPO mimetic agent eltrombopag in the restoration of bone marrow cellularity and multi-lineage haemopoiesis in patients with severe refractory aplastic anemia (37). Additionally, megakaryocyte lineage skewing of pre-progenitor cells derived from *Mpl*^*PF4cre/PF4cre*^ mice suggests excessive TPO stimulation may also act during these earlier stages in the hematopoietic hierarchy for megakaryocyte lineage specification. In more differentiated progenitors, identification of Mpl receptor expression on bipotential progenitor populations that were expanded in *Mpl*^*PF4cre/PF4cre*^ mice and were associated with megakaryocytosis and a relative paucity of erythroid precursors, confirms our previous findings (23). Thus, bipotenital erythro-megakaryocytic progenitors express Mpl and are critical effector cells of TPO-dependent thrombopoiesis that are driven to differentiate into megakaryocytes in response to TPO, and which under conditions of excessive TPO stimulation, are skewed toward megakaryocytopoiesis at the expense of erythropoiesis. Despite increased number of diverse myeloid progenitor cells in *Mpl*^*PF4cre/PF4cre*^ mice, the number of circulating white blood cells was not consistently elevated, suggesting the contribution of other regulatory mechanisms in control of mature granulocyte and macrophage number.

Finally, the demonstration that the chronic TPO stimulation gene expression signature in LSK cells of *Mpl*^*PF4cre/PF4cre*^ mice was also evident in the gene expression changes in human bone marrow CD34^+^ cells from patients with essential thrombocythemia, provides an important potential explanation for the observation that abnormally low expression of MPL in platelets and megakaryocytes of human patients with myeloproliferative disorders is associated with thrombocytosis (31, 32). Our data support a model that such disorders may be in part underpinned by insufficient MPL mass within the platelet/megakaryocyte pool resulting in increased TPO stimulation of the MPL expressing stem and progenitor cells similar to that observed in *Mpl*^*PF4cre/PF4cre*^ mice.

## Materials and Methods

Mice were analysed at age 7-12 weeks. *TPO*^*Tg*^ (3) and *Mpl*^−/−^ (24) mice have been previously described. See “SI Appendix, *SI Materials and Methods”* for generation of *Mpl*^*fl/fl*^, *Mpl*^*Delcre/Delcre*^and *Mpl*^*PF4cre/PF4cre*^ mice. Experiments were performed using procedures approved by The Walter and Eliza Hall Institute of Medical Research Animal Ethics Committee. Haematology, histology, clonal analysis of bone marrow cells semisolid agar cultures, liquid cultures of progenitor cells, flow cytometry, megakaryocyte analysis, Western blot analysis, platelet life span, serum TPO measurement, reverse transcription PCR, statistical analysis and bioinformatic analysis are described in “SI Appendix, *SI Materials and Methods*”. Microarray data are available at Array Express (www.ebi.ac.uk/arrayexpress/) under accession number E-MTAB-2389.

## Acknowledgements

We thank Janelle Lochland, Jason Corbin, Emilia Simankowicz, Melanie Howell, Lauren Wilkins and Keti Stoev for skilled assistance. This work was supported by Program Grants (1016647, 490037), Fellowships (WSA, 575501), and Independent Research Institutes Infrastructure Support Scheme Grant (361646) from the Australian National Health and Medical Research Council, the Carden Fellowship (DM) of the Cancer Council, Victoria, the Cure Cancer Australia/Leukaemia Foundation Australia Post Doctoral Fellowship and Lions Fellowship, Cancer Council of Victoria (APN), the Australian Cancer Research Fund and Victorian State Government Operational Infrastructure Support.

## References

1. Kuter DJ (2013) Thrombopoietin Mimetics. Platelets, ed Michelson AD (Academic Press, London), pp 1217–1242.

2. Qian S, Fu F, Li W, Chen Q, & de Sauvage FJ (1998) Primary role of the liver in thrombopoietin production shown by tissue-specific knockout. Blood 92(6):2189–2191.

3. de Graaf CA, et al. (2010) Regulation of hematopoietic stem cells by their mature progeny. Proceedings of the National Academy of Sciences of the United States of America 107(50):21689–21694.

4. Lannutti BJ, Epp A, Roy J, Chen J, & Josephson NC (2009) Incomplete restoration of Mpl expression in the mpl-/- mouse produces partial correction of the stem cell-repopulating defect and paradoxical thrombocytosis. Blood 113(8):1778–1785.

5. Tiedt R, et al. (2009) Pronounced thrombocytosis in transgenic mice expressing reduced levels of Mpl in platelets and terminally differentiated megakaryocytes. Blood 113(8): 1768–1777.

6. Emmons RV, et al. (1996) Human thrombopoietin levels are high when thrombocytopenia is due to megakaryocyte deficiency and low when due to increased platelet destruction. Blood 87(10):4068–4071.

7. Griesshammer M, et al. (1998) High levels of thrombopoietin in sera of patients with essential thrombocythemia: cause or consequence of abnormal platelet production? Annals of hematology 77(5):211–215.

8. Nurden AT, Viallard JF, & Nurden P (2009) New-generation drugs that stimulate platelet production in chronic immune thrombocytopenic purpura. Lancet 373(9674):1562–1569.

9. Uppenkamp M, Makarova E, Petrasch S, & Brittinger G (1998) Thrombopoietin serum concentration in patients with reactive and myeloproliferative thrombocytosis. Annals of hematology 77(5):217–223.

10. Arnold JT, et al. (1997) A single injection of pegylated murine megakaryocyte growth and development factor (MGDF) into mice is sufficient to produce a profound stimulation of megakaryocyte frequency, size, and ploidization. Blood 89(3):823–833.

11. Broudy VC, Lin NL, & Kaushansky K (1995) Thrombopoietin (c-mpl ligand) acts synergistically with erythropoietin, stem cell factor, and interleukin-11 to enhance murine megakaryocyte colony growth and increases megakaryocyte ploidy in vitro. Blood 85(7):1719–1726.

12. de Sauvage FJ, et al. (1996) Physiological regulation of early and late stages of megakaryocytopoiesis by thrombopoietin. The Journal of experimental medicine 183(2):651–656.

13. Drachman JG, Sabath DF, Fox NE, & Kaushansky K (1997) Thrombopoietin signal transduction in purified murine megakaryocytes. Blood 89(2):483–492.

14. Leven RM, Clark B, & Tablin F (1997) Effect of recombinant interleukin-6 and thrombopoietin on isolated guinea pig bone marrow megakaryocyte protein phosphorylation and proplatelet formation. Blood Cells Mol Dis 23(2):252–268.

15. Tajika K, Nakamura H, Nakayama K, & Dan K (2000) Thrombopoietin can influence mature megakaryocytes to undergo further nuclear and cytoplasmic maturation. Exp Hematol 28(2):203–209.

16. Zucker-Franklin D & Kaushansky K (1996) Effect of thrombopoietin on the development of megakaryocytes and platelets: an ultrastructural analysis. Blood 88(5):1632–1638.

17. Bunting S, et al. (1997) Normal platelets and megakaryocytes are produced in vivo in the absence of thrombopoietin. Blood 90(9):3423–3429.

18. Choi ES, et al. (1996) The role of megakaryocyte growth and development factor in terminal stages of thrombopoiesis. British journal of haematology 95(2):227–233.

19. Vitrat N, et al. (1998) Compared effects of Mpl ligand and other cytokines on human MK differentiation. Stem Cells 16 Suppl 2:37–51.

20. Tiedt R, Schomber T, Hao-Shen H, & Skoda RC (2007) Pf4-Cre transgenic mice allow the generation of lineage-restricted gene knockouts for studying megakaryocyte and platelet function in vivo. Blood 109(4):1503–1506.

21. Calaminus SD, et al. (2012) Lineage tracing of Pf4-Cre marks hematopoietic stem cells and their progeny. PloS one 7(12):e51361.

22. Srinivas S, et al. (2001) Cre reporter strains produced by targeted insertion of EYFP and ECFP into the ROSA26 locus. BMC developmental biology 1:4.

23. Ng AP, et al. (2012) Characterization of thrombopoietin (TPO)-responsive progenitor cells in adult mouse bone marrow with in vivo megakaryocyte and erythroid potential. Proc Natl Acad Sci USA 109(7):2364–2369.

24. Alexander WS, Roberts AW, Nicola NA, Li R, & Metcalf D (1996) Deficiencies in progenitor cells of multiple hematopoietic lineages and defective megakaryocytopoiesis in mice lacking the thrombopoietic receptor c-Mpl. Blood 87(6):2162–2170.

25. Rodriguez CI, et al. (2000) High-efficiency deleter mice show that FLPe is an alternative to Cre-loxP. Nature genetics 25(2):139–140.

26. Metcalf D, et al. (2008) Two distinct types of murine blast colony-forming cells are multipotential hematopoietic precursors. Proceedings of the National Academy of Sciences of the United States of America 105(47):18501–18506.

27. Metcalf D, Ng A, Mifsud S, & Di Rago L (2010) Multipotential hematopoietic blast colony-forming cells exhibit delays in self-generation and lineage commitment. Proceedings of the National Academy of Sciences of the United States of America 107(37):16257–16261.

28. Metcalf D, et al. (2009) Murine hematopoietic blast colony-forming cells and their progeny have distinctive membrane marker profiles. Proceedings of the National Academy of Sciences of the United States of America 106(45):19102–19107.

29. Smyth GK (2004) Linear models and empirical bayes methods for assessing differential expression in microarray experiments. Statistical applications in genetics and molecular biology 3:Article3.

30. Wu D, et al. (2010) ROAST: rotation gene set tests for complex microarray experiments. Bioinformatics 26(17):2176–2182.

31. Horikawa Y, et al. (1997) Markedly reduced expression of platelet c-mpl receptor in essential thrombocythemia. Blood 90(10):4031–4038.

32. Harrison CN, et al. (1999) Platelet c-mpl expression is dysregulated in patients with essential thrombocythaemia but this is not of diagnostic value. British journal of haematology 107(1):139–147.

33. Catani L, et al. (2009) Molecular profile of CD34+ stem/progenitor cells according to JAK2V617F mutation status in essential thrombocythemia. Leukemia 23(5):997–1000.

34. Kuter DJ (2007) New thrombopoietic growth factors. Blood 109(11):4607–4616.

35. Kiel MJ, et al. (2005) SLAM family receptors distinguish hematopoietic stem and progenitor cells and reveal endothelial niches for stem cells. Cell 121(7):1109–1121.

36. Mendez-Ferrer S, et al. (2010) Mesenchymal and haematopoietic stem cells form a unique bone marrow niche. Nature 466(7308):829–834.

37. Olnes MJ, et al. (2012) Eltrombopag and improved hematopoiesis in refractory aplastic anemia. The New England journal of medicine 367(1): 11–19.

